# Interplay between the epigenetic enzyme lysine (K)-specific demethylase 2B and Epstein-Barr virus infection

**DOI:** 10.1101/367094

**Authors:** Romina C. Vargas-Ayala, Antonin Jay, Hector Hernandez-Vargas, Audrey Diederichs, Alexis Robitaille, Cecilia Sirand, Maria Grazia Ceraolo, Maria Romero, Marie Pierre Cros, Cyrille Cuenin, Geoffroy Durand, Florence Le Calvez-Kelm, Lucia Mundo, Mohamed Ali Maroui, Lorenzo Leoncini, Evelyne Manet, Zdenko Herceg, Henri Gruffat, Rosita Accardi

**Affiliations:** International Agency for Research on Cancer, World Health Organization, Lyon, France; Lyon Cancer Research Center (CRCL), INSERM U1052, Centre Léon Bérard, Lyon, France; Tumor Immunology Unit, Division of Immunology, Transplantation and Infectious Diseases, IRCCS San Raffaele Scientific Institute, Milan, Italy; Department of Medical Biotechnology, Section of Pathology, University of Siena, Siena, Italy; CIRI, Centre International de Recherche en Infectiologie, (Oncogenic Herpesviruses Team), Univ Lyon; Inserm, U1111; Université Claude Bernard Lyon 1; CNRS, UMR5308; ENS de Lyon; F-69007, Lyon, France

**Keywords:** EBV, epigenetic, Burkitt lymphomas, KDM2B

## Abstract

Histone modifier lysine (K)-specific demethylase 2B (KDM2B) plays a role in hematopoietic cells differentiation and its expression appears to be deregulated in certain cancers of hematological and lymphoid origins. We have previously found that KDM2B gene is differentially methylated in cell lines derived from the Epstein-Barr virus (EBV) associated endemic Burkitt’s lymphomas (eBL) compared to EBV negative sporadic BL cells. However, whether KDM2B plays a role in eBL development has never been previously demonstrated. Oncogenic viruses have been shown to hijack the host cell epigenome to complete their life cycle and to promote the transformation process by perturbing cell chromatin organization. Here we investigated whether EBV would alter KDM2B levels to enable its life cycle and promote B-cells transformation. We show that infection of B-cells with EBV leads to down-regulation of KDM2B levels. We also show that LMP1, one of the main EBV transforming proteins, induces increased DNMT1 recruitment to KDM2B gene and augments its methylation. By altering KDM2B levels and performing chromatin immunoprecipitation in EBV infected B-cells, we were able to show that KDM2B is recruited to the EBV gene promoters and inhibits their expression. Furthermore, forced KDM2B expression in immortalized B-cells led to altered mRNA levels of some differentiation-related genes. Our data show that EBV deregulates KDM2B levels through an epigenetic mechanism and provide evidence for a role of KDM2B in regulating virus and host cell gene expression, warranting further investigations to assess the role of KDM2B in the process of EBV-mediated lymphomagenesis.

IMPORTANCE. In Africa, Epstein-Barr virus infection is associated with endemic Burkitt lymphoma, a pediatric cancer. The molecular events leading to its development are poorly understood compared to the sporadic Burkitt lymphoma. In a previous study, by analyzing the DNA methylation changes in endemic compared to sporadic Burkitt lymphomas cell lines, we identified several differential methylated genomic positions in proximity of genes with a potential role in cancer, among them the KDM2B gene. KDM2B encodes a histone H3 demethylase already shown to be involved in some hematological disorders. However, whether KDM2B plays a role in the development of Epstein-Barr virus-mediated lymphoma has never been investigated before. In this study we show that Epstein-Barr virus deregulates KDM2B expression and describe the underlying mechanisms. We also reveal a role of the demethylase in controlling viral and B-cells genes expression, thus highlighting a novel interaction between the virus and the cellular epigenome.

## Introduction

Epstein Barr Virus (EBV) is a human gammaherpesvirus. More than 95% of the human worldwide adult population is infected by EBV. After infection, EBV establishes a life-long latency, with often no adverse consequence for health. Despite its ubiquity, EBV infection is also associated with many human cancers, among them endemic Burkitt’s lymphoma (eBL) the most common childhood cancer in equatorial Africa (1). This malignancy has been associated with EBV infection more than 50 years ago, however to date the exact mechanism by which the virus contributes in the eBL pathogenic process is not fully understood.

Many studies have highlighted a key role of epigenetic deregulations in cell transformation and cancer development. Increasing evidence argues that different viruses may abrogate cellular defense systems by hijacking epigenetic mechanisms to deregulate host cell gene expression program and modulate their own life cycle (2, 3). Our recent study of the methylome profiles of sporadic- versus endemic-derived BL cell lines revealed EBV infection-specific pattern of methylation, with aberrant methylation in genes with a known role in lymphomagenesis, such as ID3, often found mutated in sporadic BL (sBL) (4). We therefore hypothesized that a viral-driven mechanism is responsible for modifying the epigenome of B-cells to facilitate the lymphomagenic process, circumventing the need for mutations in lymphoma driver genes. Among the genes differentially methylated in eBL compared to sBL, we identified the KDM2B gene, which encodes a histone H3 demethylase known to target specific sites such as trimethylated Lysine 4 (3me-K4) and di-methylated Lysine 36 (2me-K36). KDM2B sets up the stage for DNA methylation and gene silencing by recruitment of the polycomb-1 proteins to un-methylated CpG regions (5) and plays a key role in somatic cells reprogramming (6). It also represses the transcription of ribosomal RNA genes inhibiting cell growth and proliferation (7). KDM2B has been identified as a putative tumor suppressor by retroviral insertion analysis in mice (8). Low level of KDM2B expression has been found in aggressive brain tumors, suggesting its potential role in cancer development. Moreover, KDM2B plays a key role in hematopoietic cell development and displays opposing roles in tumors of hematopoietic and lymphoid origins (9). While high level of KDM2B expression has been observed in different hematological malignancies, its depletion from hematopoietic cells has been reported to activate cell cycle and reduce the activity of interferon and lymphoid-specific transcription factors, thereby contributing to myeloid transformation (9). However, whether KDM2B affects EBV life cycle has never been determined, nor its role in endemic BL has been assessed. Here, using in vitro EBV-infection models, we aimed to assess whether the virus can alter the expression of KDM2B by inducing its gene methylation. Finally we investigated how this event impact on EBV infection and B-cells homeostasis. Overall, our data highlight a novel crosstalk between EBV and the cellular epigenome, identify KDM2B as a master regulator of EBV gene expression in addition to B-cell gene expression, suggesting a role for EBV-mediated KDM2B deregulation in the lymphomagenic process.

## Results

### KDM2B is epigenetically silenced in EBV+ BL derived cell lines

Our comparative analysis of the whole genome methylation profiles of a set of EBV (+) and EBV (-) Burkitt’s lymphoma derived cell lines (4), led to identification of two CpGs (CpG 15695155 and 21423404) in an intragenic putative regulatory region of KDM2B (Fig 1A) highly methylated in EBV (+) BL derived cells. To validate these data we performed direct pyrosequencing on DNA extracted from 7 BL EBV (+) BL and 7 EBV (-) BL derived cell lines and confirmed that KDM2B gene is hyper-methylated at CpG 15695155 and 21423404 in EBV (+) BL cell lines compared to EBV (–) BL cell lines (Figure 1B). Next we asked whether these different patterns of methylation affect KDM2B expression level. Four EBV (+) BL and 3 EBV (-) BL samples were analyzed by immunohistochemistry for the KDM2B protein expression level. Three out of four EBV (+) samples showed a weaker staining compared to the EBV (-) BLs, suggesting that EBV infection induces a reduced expression of KDM2B protein *in vivo* (Fig 1C). Of note, the EBV (+) sample with stronger KDM2B staining, had a fewer amount of EBER expressing cells, as shown by the EBER FISH (data not shown). We next assessed whether silencing of KDM2B expression in EBV (+) BL cell lines was mediated by DNA methylation. Treating 3 EBV (-) BL and 3 EBV (+) BL cell lines with the demethylation agent 5-Aza-2’-deoxycytidine (Aza) for 48h led to a significant rescue of KDM2B expression level in EBV (+) BL, whereas no noticeable changes in KDM2B mRNA expression was observed in EBV (-) BL cells (Fig 1D). In conclusion, our data show that the specific CpG sites in the regulatory region of KDM2B are methylated in EBV (+) BL compared to EBV (-) BL cell lines confirming our previous whole methylation profiling data (4). Moreover, silencing of KDM2B expression appears to be a frequent event in endemic BLs specimens and to be mediated by DNA methylation. These data indicate that EBV may regulate KDM2B expression level by inducing methylation at a specific regulatory region of the gene.

**Figure 1.**
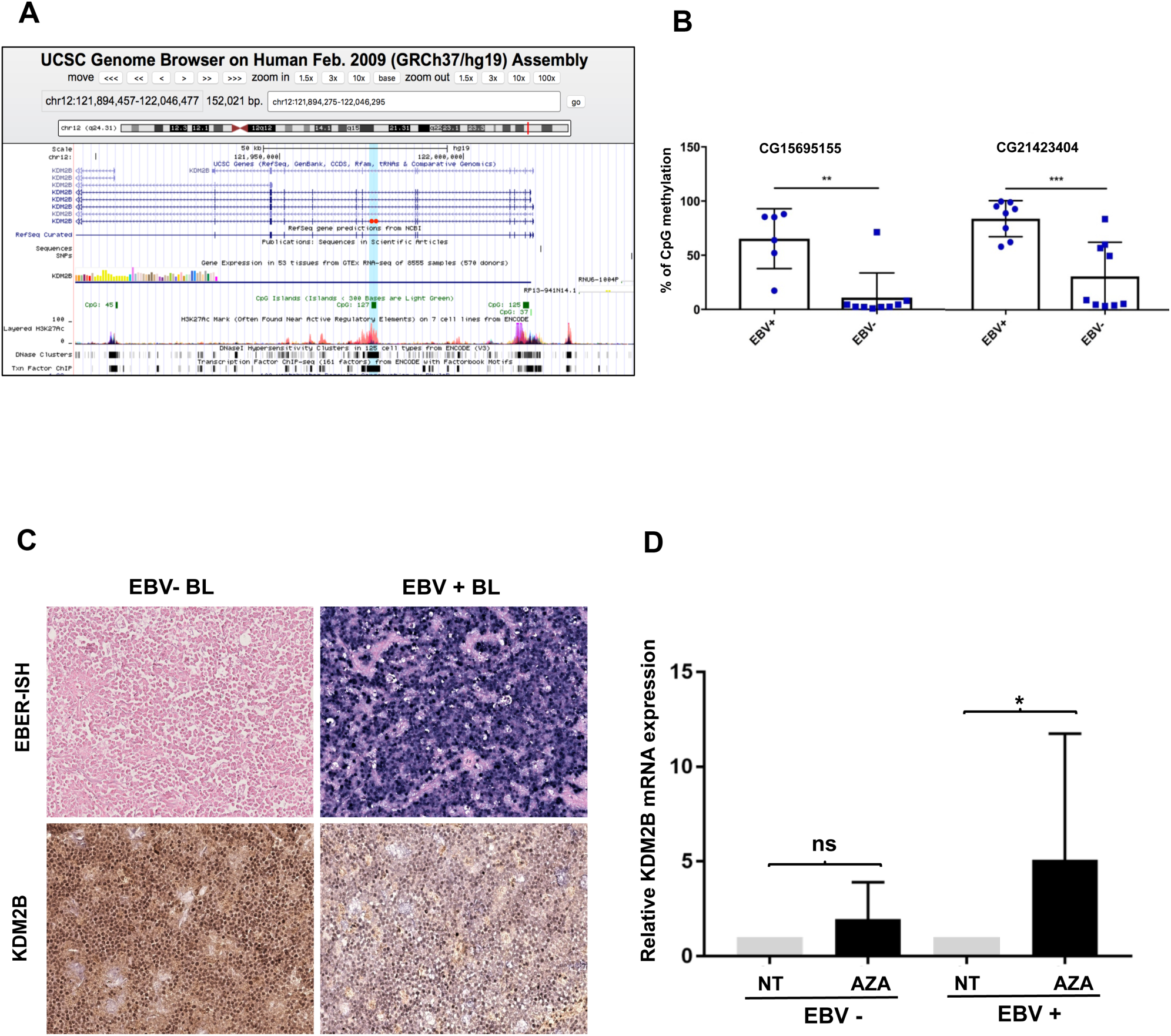
KDM2B gene is methylated and silenced in EBV+ BL cell lines and specimens. **A**. Schema of KDM2B gene (modified from UCSC genome browser). Red dots show CpG positions. **B**. The histograms show the average % of methylation of cg15695155 and cg21423404 in the DNA of 7 EBV (+) and 7 EBV (-) BL, measured by pyrosequencing (**p value<0.01, ***p value<0.001). **C**. KDM2B level in EBV (+) and EBV (-) BL samples were analyzed by immunohistochemistry. The same samples were analyzed for the expression of EBER by EBER-ISH as explained in the material and methods section. **D**. Three EBV (+) and 3 EBV (-) BL were cultured in presence of 5-Aza-2’-deoxycytidine (AZA) at the final concentration of 10µM for 48h (AZA=treated) or with DMSO (NT=untreated). Messenger RNA levels of KDM2B were analyzed by qPCR. The pooled results of three independent Aza treatment are represented in the histograms (*p value<0.05).

### Infection of B cells with EBV leads to down-regulation of KDM2B levels

To assess the ability of EBV to deregulate KDM2B expression level we infected B-cells isolated from 4 independent donors with EBV and let the cells immortalize to generate lymphoblastoid cell lines (LCLs). The expression level of KDM2B in these cell lines were then compared to that in the parental counterparts by RT-qPCR. LCL showed reduced expression level of KDM2B transcripts compared to primary B-cells (Fig 2A), highlighting a role of EBV infection in the regulation of KDM2B mRNA expression. To evaluate whether downregulation of KDM2B in LCLs compared to primary B cells was due to infection with EBV or occurred as a side effect of the immortalization process, we monitored the changes in the expression level of KDM2B at different time points during the process of *in vitro* EBV-mediated B cells immortalization. To this end, B-cells from different donors were infected with EBV. Cells were harvested at 48h, 96h and 4 weeks after infection, when LCLs were established, processed for RNA isolation and analyzed by RT-qPCR for the expression level of KDM2B. Forty-eight hours after EBV infection we could already observe a significant downregulation of KDM2B mRNA and protein levels compared to the parental primary B-cells (Fig 2B-C), indicating that silencing of KDM2B takes place early upon EBV infection and does not require B-cells immortalization to occur. To exclude that this result could be due to the different proliferation status between LCL or BL derived cell lines and primary B cells, we infected the RPMI immortalized B cells line with EBV and compared them to mock infected RPMI cells for the levels of KDM2B mRNA and protein. The results show that KDM2B mRNA and protein expression is reduced 48h upon EBV infection both at the RNA and protein levels, respectively in EBV infected RPMI compared to mock infected cells (Fig 2D and E), confirming the results obtained in EBV-infected primary B-cells.

**Figure 2.**
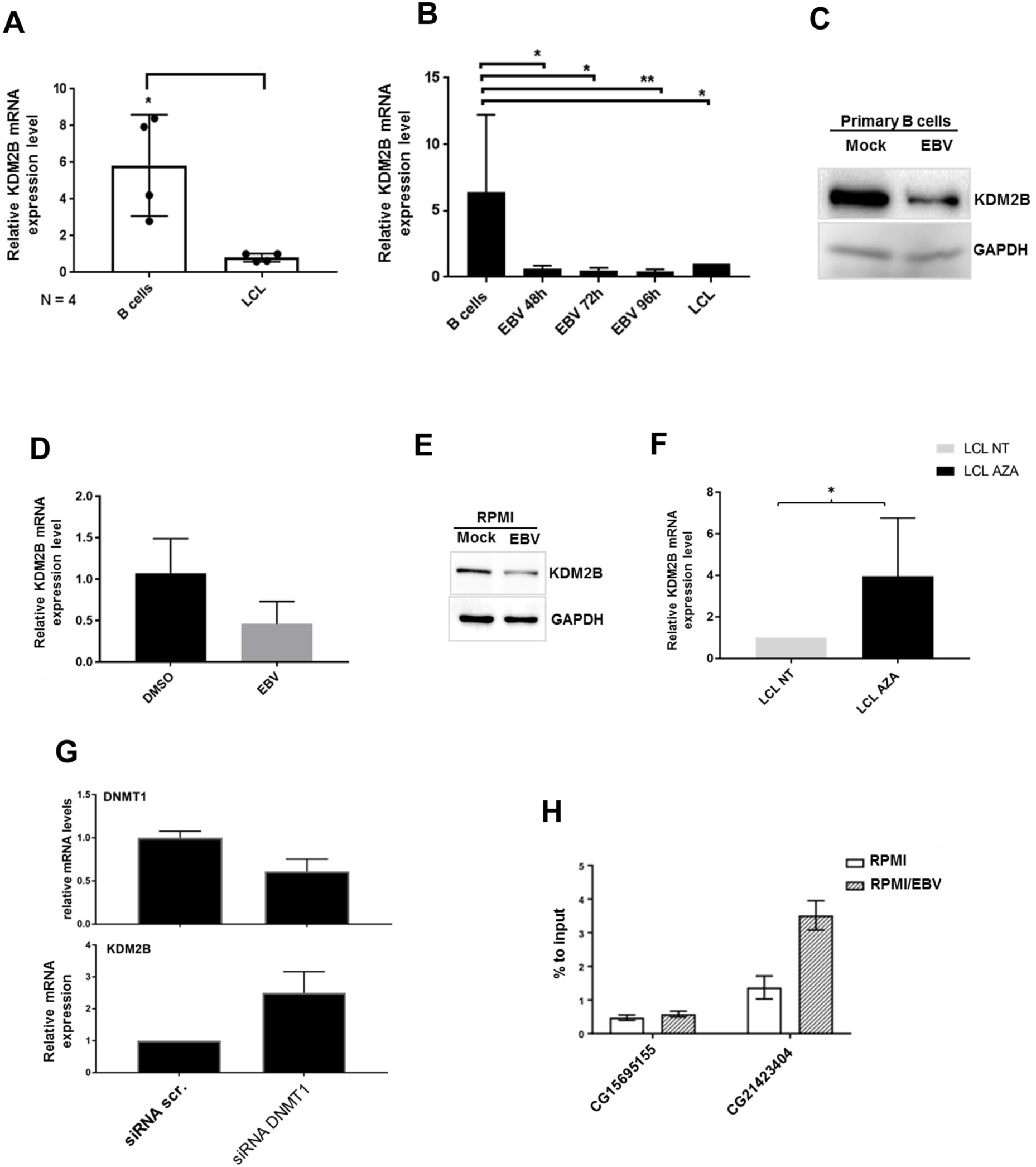
EBV-dependent silencing of KDM2B expression *in vitro*. **A**. Primary B cells from four different donors were isolated and cultured for 24-36h prior to their infection, after EBV infection, cells were kept in culture until they got immortalized (LCL).. Nucleic acids were extracted from primary and immortalized cells, total RNA retro-transcribed and analyzed by qPCR for the levels of KDM2B. The histogram shows the average KDM2B mRNA levels observed in B cells and in the corresponding LCL from four independent donors (* p value <0.05). **B-C**. Primary B cells from three independent donors were EBV infected, in part collected to make dry pellets at the indicated time points and in part let in culture to generate LCL. Cell were processed and analyzed for the levels of KDM2B mRNA by qPCR (**B)**. Total proteins were extracted by B cells and B cells at 48h post EBV infection and were analyzed by western blot for the indicated proteins (**C**). **D-E**. RPMI cells were infected with EBV or mock infected for 48h, then collected and processed for RNA/protein extraction. Total RNA was retro-transcribed and cDNA analyzed by qPCR for the levels of KDM2B (**D**). Total proteins were analyzed for the levels of the indicated protein by western blot (**E)**. **F**. LCL were cultured for 96h in presence of DMSO or 5-Aza-2’-deoxycytidine (AZA) used as in Figure 1C. Cells were then collected and the extracted mRNA were analyzed for the levels of KDM2B. **G**. LCL were transfected as explained in Methods section with stabilized DNMT1 siRNA. Four days after transfection cells were collected and analyzed for the levels of mRNA of DNMT1 (upper panel) and KDM2B (lower panel). **H** RPMI cells infected as for figure 2D-E were fixed and processed for ChIP with a DNMT1 antibody and IgG as negative control. The eluted DNA was analysed by qPCR with primers flanking the cg15695155 and cg21423404 (primer sequences described in Table 1).

To determine whether the reduced expression of KDM2B in EBV immortalized cells could be due to an increase in DNA methylation, LCLs were treated with the demethylating agent Aza and analyzed by RT-qPCR for the expression level of KDM2B. Aza-treated LCLs showed increased expression level of KDM2B mRNA, compared to their untreated counterpart, indicating that downregulation of KDM2B in EBV infected cells is mediated by DNA methylation (Fig 2F). As incorporation of Aza into the DNA impedes its methylation by DNMTs (10), we hypothesized that EBV could contribute to KDM2B silencing by increasing the recruitment of DNMT1 onto the KDM2B gene. Indeed depletion of DNMT1 in LCLs by transfection of DNMT1 siRNA led to an increase level of KDM2B mRNA expression (Fig 2G). Moreover, ChIP experiments showed an increased recruitment of DNMT1 onto the CpG site of the KDM2B gene in RPMI cells infected with EBV (Fig 2 H).

Taken together, these results indicate that EBV infection in B-cells leads to a reduced expression of KDM2B. DNMT1-mediated methylation of KDM2B gene appears to play a role in this event.

### The oncogenic viral protein LMP1 induces the silencing of KDM2B

Next we asked whether LMP1, the main EBV oncoprotein, may play a role in the deregulation of KDM2B expression. To this aim, we have generated RPMI cells stably expressing LMP1. As a negative control, the cells were transduced with the empty retroviral vector (pLNSX) (Fig. 3A). As revealed by the RT-qPCR analysis, compared to RPMI-pLXSN, RPMI-LMP1 cells display lower expression levels of KDM2B mRNA and protein (Fig 3B-C). Thus, LMP1 appears to play a role in EBV-mediated downregulation of KDM2B expression. In addition, treating RPMI-LMP1 cells with Aza led to a rescue of KDM2B RNA levels (Fig 3D). By contrast, no change in KDM2B mRNA expression level was observed when treating RPMI-pLXSN cells with the DNA demethylating agent. This was confirmed by pyrosequencing of the two CpG sites that were previously identified as differentially methylated in EBV (+) BL cell lines (CpG 21423404 and CpG15695255). RPMI-LMP1 cells have higher levels of DNA methylation at both CpG when compared to RPMI-pLXSN, even though the difference is higher for the CpG 15695255 (Fig 3E).

**Figure 3.**
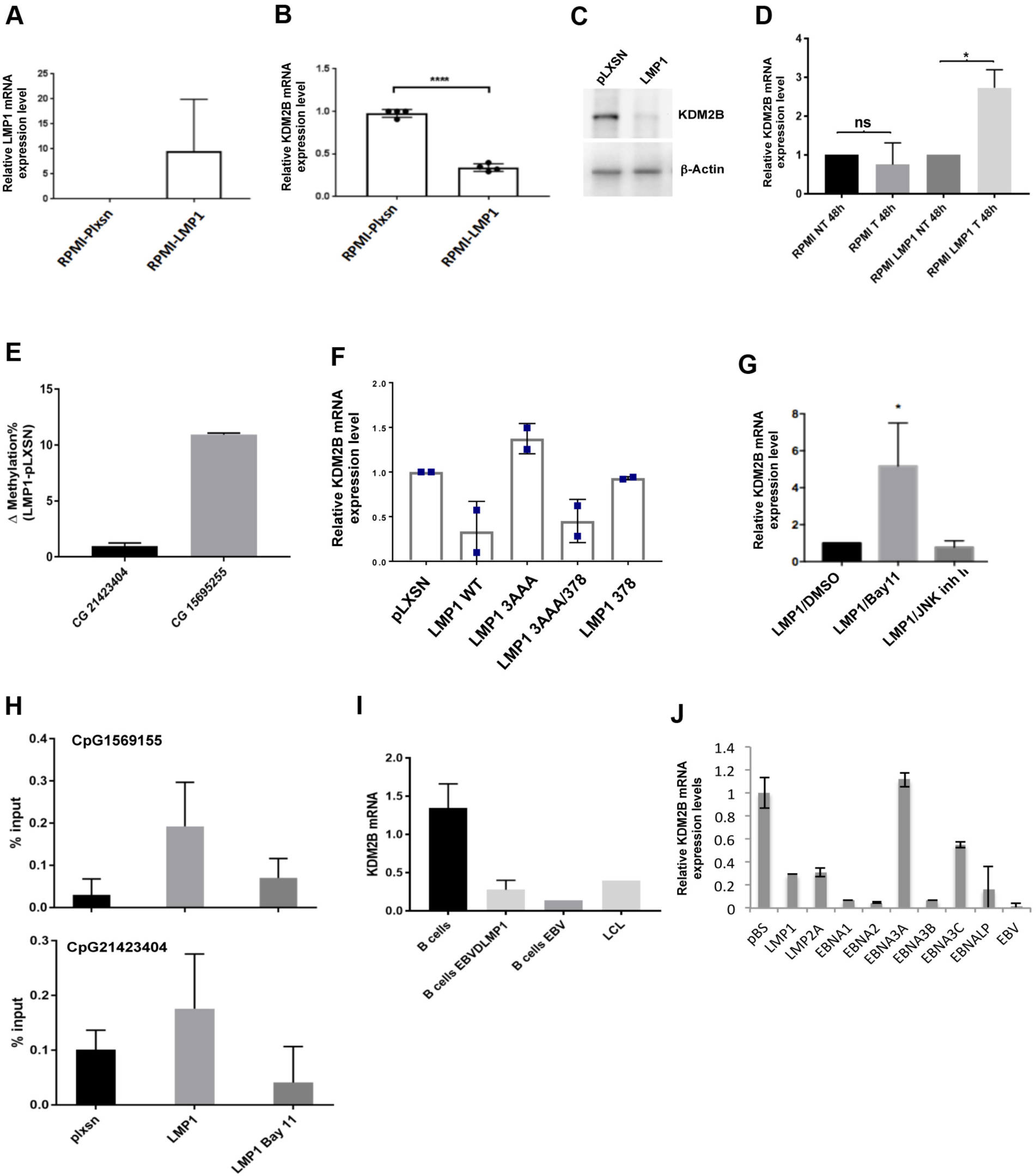
LMP1-mediated downregulation of KDM2B. **A-C** RPMI cells were stably transduced with pLXSN (pLXSN) or with pLXSN-LMP1 WT (LMP1). Cells were collected and processed for RNA/DNA and protein extraction. cDNA samples were interrogated by qPCR for the levels of LMP1 (**A**) and KDM2B transcript (****p value <0.0001) (**B**) or protein levels (**C**). **D**. RPMI or RPMI-LMP1 cells were cultured in presence of AZA (T) or in DMSO (NT) for 48h and the mRNA levels of KDM2B were analyzed by RT-qPCR. **E**. RPMI or RPMI-LMP1 cells DNA samples were analysed for the levels of methylation of KDM2B genes at the positions cg15695155 and cg21423404, measure by pyrosequencing. The difference of methylation levels between RPMI-LMP1 and pLXSN are shown in the histogram that represent the average of two experiments. **F**. RPMI cells were stably transduced with pLXSN (pLXSN) or with pLXSN-LMP1 WT (LMP1) or mutated (3-AAA, 378 and 3-AAA/378). Cells were collected and processed for total RNA extraction. cDNA samples were interrogated by qPCR for the levels of KDM2B. **G**. RPMI-LMP1 cells were treated for 2h with Bay11 (10µM), or with JNK inhibitor II (10 µM). cDNA samples were analyzed by qPCR for the mRNA levels of KDM2B (*p value <0.05). **H**. RPMI pLXSN and RPMI-LMP1 cells, the latter, treated or not with Bay11 (10µM) for 2h, were used to perform ChIP with DNMT1 and IgG antibodies. The eluted DNA was analyzed by qPCR with primers designed to surround the two CpG of interest (cg15695155 and cg21423404). The histograms show the average % of recruitment of DNMT1 in the different conditions, in two independent experiments. **I**. B-cells from different donors were infected with EBVWT or EBV_Delta LMP1_ genomes and collected at 48h from the infection. EBV WT infected B-cells were left in culture until they generate LCLs. Retro-transcribed RNA samples were analysed by qPCR for the levels of KDM2B. **J**. Louckes cells were transfected with different constructs carrying EBV genes. 48h post transfection cells were processed and analyzed for the levels of KDM2B by RT-qPCR.

LMP1 activates respectively NF-kB and JNK pathway through its CTAR1 or CTAR2 domains. Therefore, to gain insights into the mechanism by which LMP1 deregulates KDM2B expression we generated RPMI cells expressing LMP1 mutants, harboring mutations on CTAR1 (mutant 3AAA), CTAR2 (mutant 378) or on both CTARs (mutant 3AAA/378) and having therefore hampered ability to activate NFκB or JNK, or both pathways. We then compared the ability of the different LMP1 mutants to deregulate KDM2B. As shown in Figure 3F, mutant 378 retained the ability to downregulate the KDM2B mRNA level with a similar efficiency as WT LMP1. By contrast, the 3AAA mutant, as well as the double mutant, was unable to downregulate KDM2B mRNA levels (Fig. 3F), indicating that NFκB activation pathway induced by LMP1 is an important event in the regulation process of KDM2B expression. To further evaluate the impact of NFκB and JNK pathway on LMP1-mediated KDM2B deregulation, we treated RPMI-LMP1 cells with specific chemical inhibitors, BAY 11-7082 and JNK inhibitor II, respectively. Treating RPMI-LMP1 cells with the NFκB inhibitor BAY 11-7082, but not with JNKII, led to a rescue of KDM2B mRNA levels (Fig 3G), confirming that inhibition of KDM2B expression by LMP1 is mediated by the NFκB pathway.

Previous studies showed that LMPs activate DNMTs and induce their recruitment onto the promoters of cancer related genes (11-13). In line with these findings, our ChIP experiments showed an increase in the recruitment of DNMT1 onto the KDM2B gene in LMP1 expressing cells compared to the control cells (Fig 3H). Treating RPMI-LMP1 cells with BAY 11-7082 reduced significantly the amount of DNMT1 recruited into the KDM2B gene. Taken together, these data indicate that EBV induces silencing of KDM2B expression mainly *via* the ability of its main transforming protein LMP1, to activate the NF-kB signaling pathway. Nevertheless, infection of primary B-cells with a recombinant EBV lacking the LMP1 gene (EBVΔLMP1) still led to a decreased expression of KDM2B mRNA (Fig 3I), indicating that in addition to LMP1 other EBV proteins could be involved in KDM2B deregulation. Transfecting Louckes cells with different constructs expressing a panel of EBV genes (Fig 3J) confirmed that the virus may use alternative mechanisms to inhibit KDM2B expression, further indicating that this event may be important for the virus.

### KDM2B regulates viral genes expression in EBV infected B cells

To evaluate the biological relevance of EBV-mediated KDM2B expression downregulation we reasoned that the epigenetic enzyme could regulate the EBV transcription similar to what was observed for other histones modifiers and chromatin interacting proteins (e.g. EZH2, CTCF, KMT5B etc.) (14-16). To assess whether ectopic expression of KDM2B in B-cells could have an impact on EBV infection, we transfected increasing concentrations of a HA-KDM2B construct in Louckes cells, an EBV (-) BL derived cell lines. One day post-transfection cells were infected with EBV/GFP and monitored for the infection efficiency 24h later. Figure 4 A shows efficient overexpression of ectopic HA-KDM2B at 24h post transfection (Fig. 4A). Infection efficiency of Louckes cells transfected with KDM2B expression vector and the control empty vector (pCDNA) was indistinguishable as revealed by similar percentages of GFP-positive cells in the two populations analyzed by FACS (Fig 4C). This was confirmed by the Taqman PCR showing that increased KDM2B expression level did not alter viral genome copy number (Fig. 4B). By contrast, the mean green fluorescence intensity (MFI) decreased in presence of enhanced KDM2B expression level (Fig. 4D), indicating that KDM2B could affect viral gene expression. Indeed, RT-qPCR analysis of the expression level of different EBV transcripts (LMP1, EBNA1 and BZLF1) at 24h post-infection showed a significant and dose dependent reduction of their mRNA levels in the presence of increasing amount of KDM2B (Fig. 4E).

**Figure 4.**
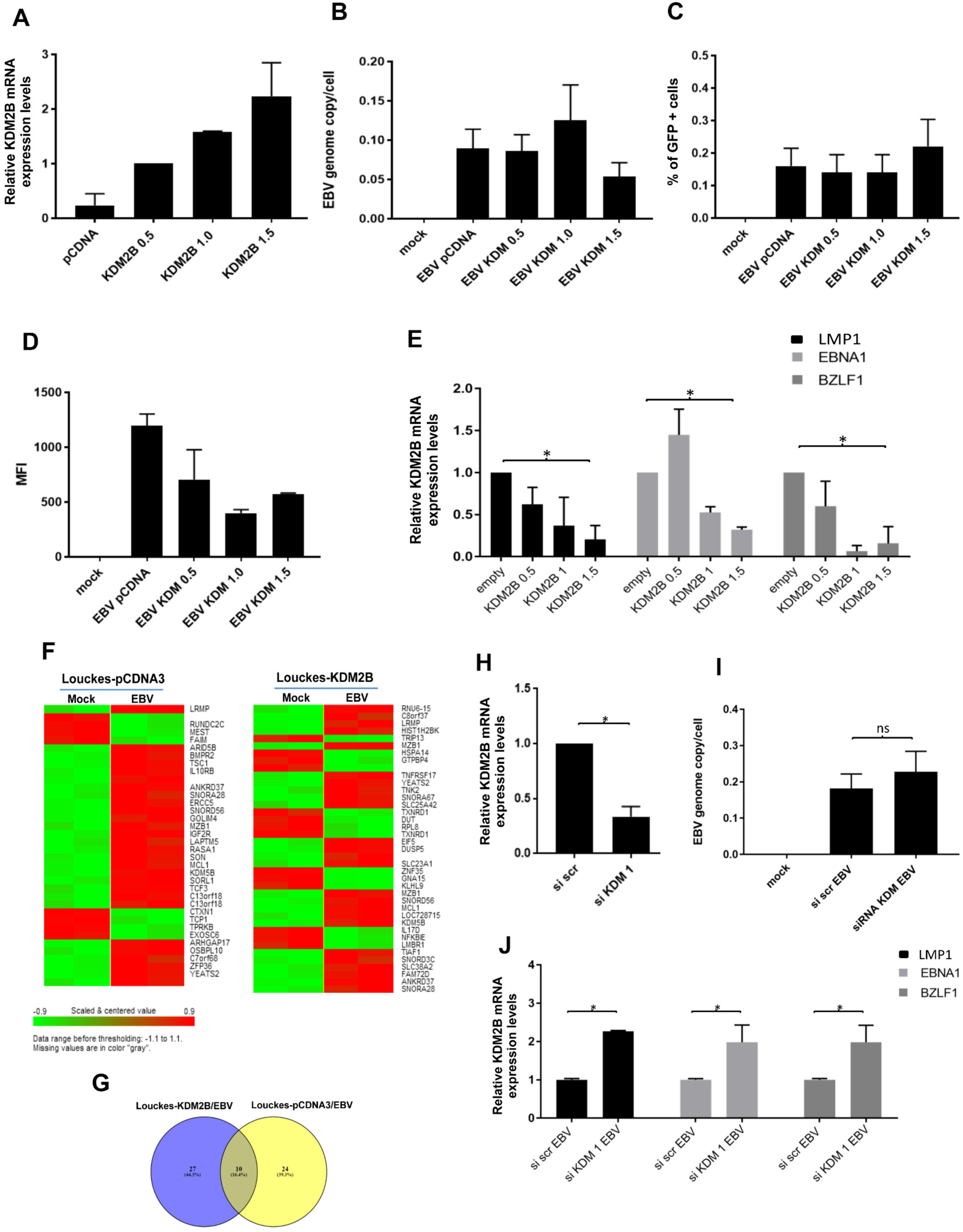
KDM2B regulates EBV gene expression in early stage of the infection. **A-G**. Louckes cells were transiently transfected with growing concentrations of KDM2B, collected at 24h post transfection and processed for RNA extraction to assess ectopic KDM2B levels by qPCR (**A**). Cells were then EBV infected and 24h after infection they were collected and processed for FACS analysis and for RNA/DNA extraction. DNA samples were used to measure EBV genome copy number by Taqman PCR (**B**). Live cells were analyzed for the percentage of GFP+ cells and for the mean fluorescence intensity signal (MFI) by FACS. (**C-D**). cDNA samples were analyzed for the expression levels of EBV early and late genes by qPCR. The values showed in the histogram are the average of three independent experiments (* p value <0.05) (**E**). **F**. RNA samples were processed for RNA expression profiling. The heat map on the left side shows the genes differentially regulated in mock infected Louckes when compared to EBV infected Loucks, while the heat map on the right side compares genes deregulated in Louckes overexpressing KDM2B which are mock or EBV infected. **G**. Venny diagram compares the two gene lists identified in the comparison described in Fig 4F. **H-J**. Louckes cells were transfected with KDM2B siRNA and scrambled siRNA as control. 24 hours upon transfection cells were in part collected and analyzed for the levels of KDM2B mRNA by qPCR (**H**), and in part infected with EBV. 24h post infection cells were collected and processed for RNA/DNA extraction. While the EBV genome copies per cells was determined by Taqman PCR on the DNA template (**I)**. QPCR on the cDNA samples allowed assessing the mRNA expression levels of different viral genes **(J)**.

Furthermore, RNA expression analysis showed that gene expression profile of EBV infected Louckes cells, changed in presence of high ectopic level of KDM2B (Fig. 4F-G). Indeed, in line with reduced expression level of EBV genes transcripts, the deregulated expression of host genes known for being altered during EBV infection, such as IL10RB (17) or RUND2C, was attenuated or completely lost in the presence of increased KDM2B expression level (Fig. 4F). Taken together, these data indicate that KDM2B plays a role in regulating the expression of EBV genes.

We next depleted KDM2B in Louckes cells by transfecting a siRNA targeting regions within KDM2B open reading frame. 24h after transfection part of the cells were collected to extract total RNAs. RT-qPCR analysis showed that the level of KDM2B mRNA was efficiently down-regulated in cells transfected with the siRNA compared to the cells transfected with a scrambled siRNA control (Fig. 4H). Cells were then infected with EBV/GFP and monitored for the infection efficiency, as above. At 24h post infection, neither the percentage of GFP positive cells nor the genome copy number changed significantly between the control cells (si-scrambled) and the cells transduced with the siRNA directed against KDM2B (Fig. 4I and data not shown), indicating that loss of KDM2B does not affect the efficiency of infection. However efficient depletion of KDM2B by si-KDM2B led to a significant increase of EBV transcripts 24h after infection (Fig. 4J). This results indicates that reduced levels of KDM2B mRNA promotes the expression of the viral genes, further confirming the ability of KDM2B to repress EBV gene expression at early stages of infection.

### KDM2B inhibits viral gene expression in latently infected EBV immortalized B cells

To test whether the activity of KDM2B in regulating EBV gene expression was required for the maintenance of the virus latency, similarly to what has been reported for other epigenetic enzymes (14), we aimed to overexpress KDM2B ectopically and examine its impact on viral gene expression in EBV immortalized cells. LCLs display detectable, even though lower, KDM2B mRNA and protein level when compared to primary B-cells (Fig 2 A-B and data not shown). Therefore KDM2B was overexpressed in LCLs by ectopic expression of a HA-KDM2B. Forty-eight hours upon transfection, LCLs were collected and processed for total protein and DNA/RNA extraction. Western blot analysis showed increased KDM2B protein level (Fig. 5A). Efficient expression of ectopic KDM2B was also confirmed by RT-qPCR (Fig. 5B). LCLs overexpressing KDM2B carried a similar number of EBV genome copies compared to LCL transfected with the control pCDNA vector (Fig. 5C). We then analyzed the mRNA expression level of different EBV transcripts by RT-qPCR. Ectopic KDM2B expression led to a significant reduction of the mRNA expression level of all the analyzed EBV genes (Fig. 5D). By contrast, depletion of KDM2B from LCL by transfecting KDM2B siRNA (Fig. 5E-F) led to a significant increase in the expression of EBV genes, compared to the expression level of the same genes in siRNA scrambled transfected LCL (Fig. 5H). Removal of KDM2B did not significantly affect EBV genome copy number (Fig. 5G). Taken together, these data indicate that KDM2B controls viral gene expression in latently infected cells.

**Figure 5.**
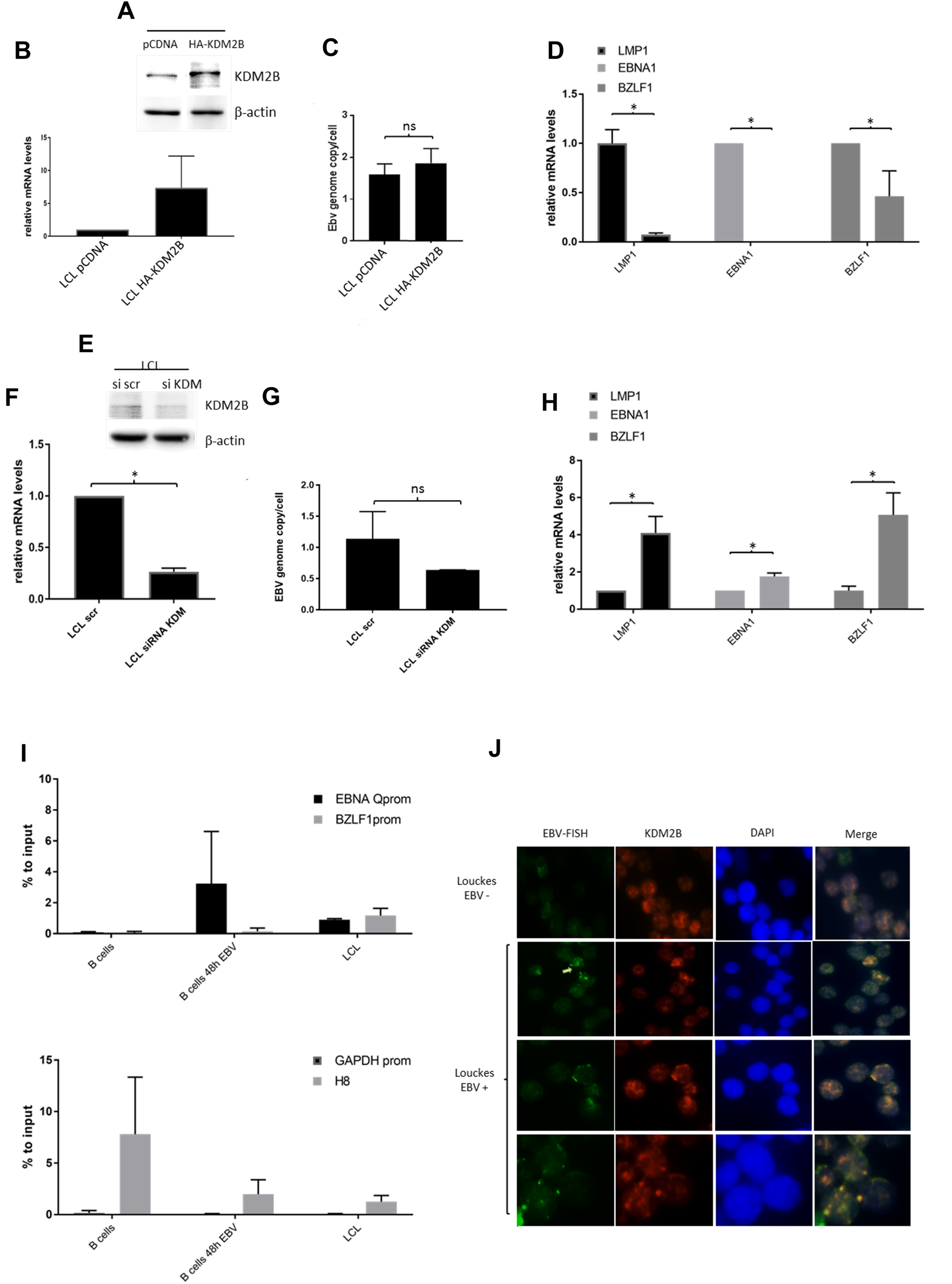
KDM2B binds to and regulates EBV gene expression in latently infected cells. **A-D**. LCLs cells were transfected with 1.5 ug of KDM2B pCDNA or with pCDNA as an empty control. Cells were collected 24h after transfection and processed for RNA/DNA and total protein extraction. Total protein extracts were analysed by western blot for the indicated proteins (**A**). KDM2B mRNA levels were assessed by qPCR (**B**). DNA samples were analyzed by Taqman PCR to assess the number of EBV genome copies per cell (**C**) The mRNA expression levels of EBV early and late genes were assessed by qPCR (**D**). **E-H** LCL transfected with KDM2B siRNA and scr siRNA as control were collected at 48h post transfection, processed and analysed for the levels of KDM2B protein (**E**) and transcript (**F**). DNA and cDNA samples were analysed respectively for the EBV DNA copy number by Taqman PCR (**G**) and for the expression levels of the EBV genes by qPCR (**H**). **I**. Primary B cells were infected with EBV for 48h or let immortalized until they generate LCL. Not infected B cells, B cells EBV infected for 48 and LCLs were formaldehyde fixed and processed for ChIP using a KDM2B antibody. The eluted DNA was analyzed by qPCR with primers designed on different EBV gene promoters. As negative controls the promoter of the GAPDH promoter was also amplified, the recruitment of KDM2B to its known cellular gene target, H8 was also assessed. **J**. Louckes cells infected or not with EBV (no GFP) were fixed on glass slides and processed for immuno-FISH. EBV DNA was detected by FISH and KDM2B by Immunofluorescence. EBV/KDM2B overlapping signals are shown in the merge fields as yellow dots.

We next asked whether KDM2B could directly bind to EBV gene promoters to regulate their expression. To this aim, purified primary B-cells, infected or mock infected with EBV for 48h, as well as the corresponding LCLs, were formaldehyde-fixed and processed for KDM2B Chromatin Immunoprecipitation (ChIP) (Fig 5I). ChIP analysis show that KDM2B can be detected on some EBV gene promoters at 48h post infection, as well as in EBV-immortalized B cells. As expected, we did not detect KDM2B recruitment on the promoter of the house keeping gene GAPDH, however we observed an efficient KDM2B binding to a sequences located downstream of the transcription start site of the rDNA repeated units 8 (H8), that was previously reported to be targeted by the epigenetic enzyme (18). This data indicates that KDM2B is directly recruited on the EBV gene promoters. To further assess the ability of KDM2B to be recruited onto the EBV genome, B cells were EBV infected or not, collected and processed for Immuno-FISH experiments. EBV DNA molecules were detected by fluorescence in situ hybridization with a labeled probe directed against the BamHI W EBV genomic repeated region; KDM2B was concomitantly detected by immunofluorescence by using an anti-KDM2B antibody. In EBV infected B-cells, KDM2B patches partially overlapped or were found in close proximity with the viral DNA (Fig 5J), suggesting that the epigenetic enzyme is recruited into or close to the viral genome early upon infection. Taken together, our data demonstrate a role of KDM2B in controlling EBV gene expression.

### KDM2B mRNA level deregulation in B cells alters their expression profile

KDM2B has been shown to play a role in differentiation, in cell growth and proliferation of hematopoietic cells (9). We then asked whether altered KDM2B expression could also have an impact on the B-cell phenotype, in our experimental models. Louckes and LCLs overexpressing KDM2B did not show altered proliferation ability, nor did they show altered cell cycle or apoptotic profiles (data not shown). Therefore, to get further insight on the impact of KDM2B deregulation on B-cells, we performed RNA expression chip array in Louckes cells expressing increasing levels of KDM2B. Cellular genes whose expression was significantly altered in the presence of increased expression level of KDM2B (Fig 6A) were identified by bioinformatics analysis. In line with its known function as demethylase of H3 K4, KDM2B altered gene set were enriched of H3 K4-me1 (Fig 6B). Among the genes significantly deregulated in the presence of altered KDM2B expression level, some played a role in immunity (Fig 6C). Our analysis revealed that KDM2B overexpression was associated with deregulated expression of genes involved in the TNFR2 pathway (Fig 6C). Interestingly, this pathway is important for the transition of B-cells from the germinative centre to resting memory B-cells (19). Moreover, TNFR2 pathway mediates specific TNF effects and is an important mediator of cell antiviral response (9). Deregulated expression of genes within this pathway were confirmed and validated by RT-qPCR (Fig 6D). Taken together, these data indicate that KDM2B plays a role in B-cells differentiation and regulation of the TNF pathway, often altered during the lymphomagenic process.

**Figure 6.**
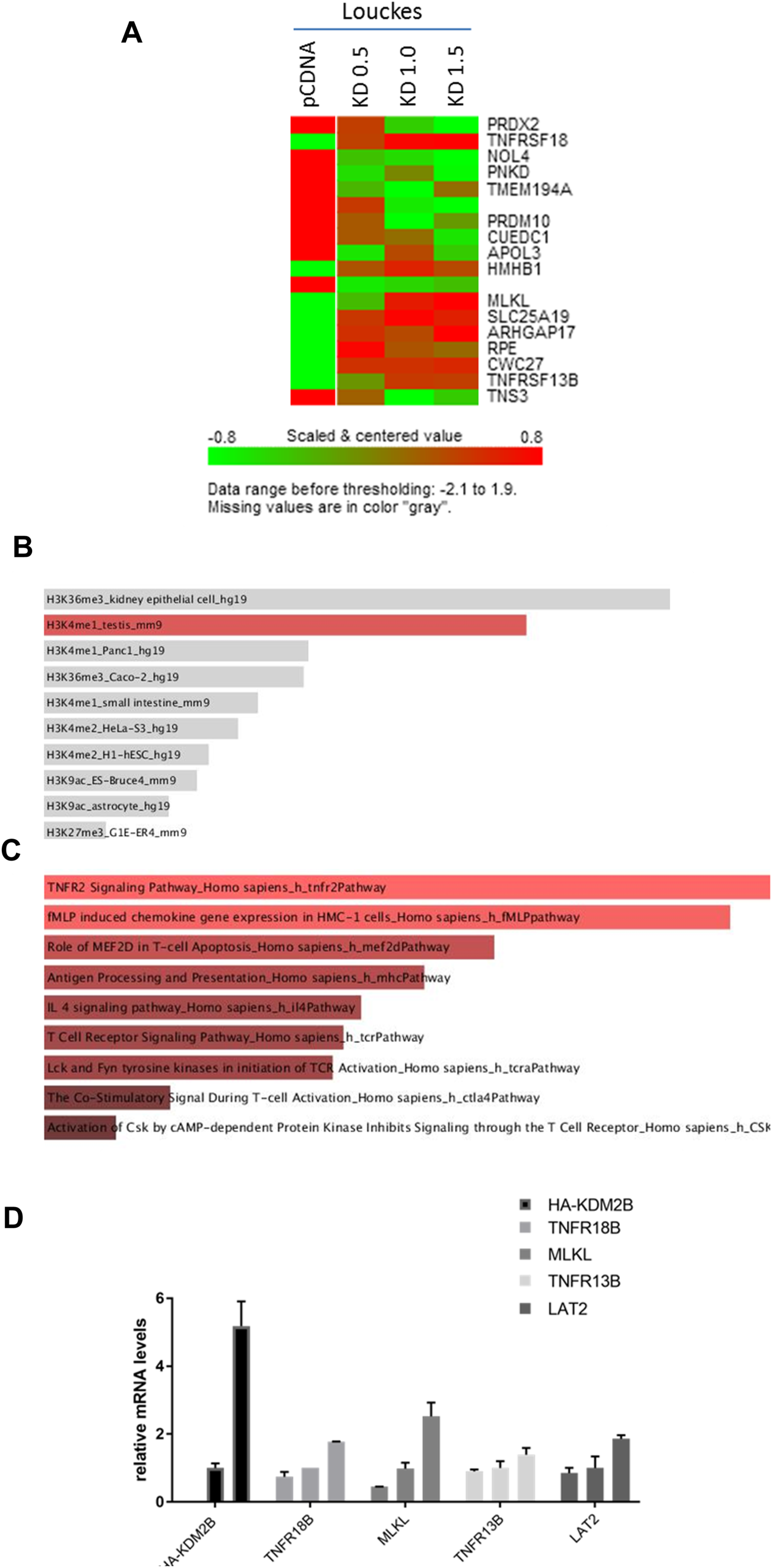
KDM2B deregulation affects cellular gene transcription. **A**. Louckes cells generated as explained in the legend of Fig 4A, were collected and processed for RNA extraction and RNA expression profiling, as described in Methods section. The differentially expression analysis was conducted using the BRB array tool**. B-C**. genes differentially regulated in Louckes overexpressing KDM2B compared to Louckes transfected with an empty pCDNA were analysed for their enrichment in specific pathways by using Enrichr web tool. Enrichment results are shown for the indicated databases (BioCarta 2016 and Epigenomics Roadmap HM ChIP-seq). **D**. A set of genes deregulated identified in the array were validated by qPCR.

## Discussion

In this study we show that a regulatory region within the gene encoding for KDM2B is more methylated in EBV (+) BL compared to EBV (-) BL, confirming data from our previous study aiming to characterize the whole epigenetic profile of a set of BL derived cell lines of endemic or sporadic origins. This region is found methylated in certain cancer derived cell lines (ENCODE DNA Methylation Tracks; CpG Methylation by Methyl 450K Bead Arrays from ENCODE/HAIB; UCSC genome browser). Furthermore, KDM2B levels appear to be deregulated in cancer of hematological origins (9). We therefore asked whether increased methylation of KDM2B gene in eBL was mediated by EBV to deregulate the histones modifier intracellular levels and promote EBV-mediated lymphomagenesis. Indeed, KDMs have been shown to have altered expression in cancer (20). For instance, a KDM2B paralogue, KDM2A, behaves as a tumor suppressor in hematopoietic stem cells, in which it antagonizes MLL-associated leukemogenesis, by erasing H3K36me2 marks (21). Similarly, altered levels of KDM2B could modify the chromatin structure and the pattern of expression of B cells and favor their transformation.

Endemic BL specimens showed low KDM2B protein level underscoring a potential important role of KDM2B downregulation in the lymphomagenic process *in vivo*. Treating eBL cell lines with a chemical blocking DNA methylation led to a rescue of KDM2B mRNA level, indicating that DNA methylation contributes to silence KDM2B expression in these cells. However the same treatment left the level of KDM2B transcript in EBV negative sporadic BL derived cell lines unchanged. These events are similar to what we have previously described for another gene, ID3 (4). This gene, which plays a key role in lymphomagenesis and which is often found mutated in the sporadic BL variant (22, 23), was found silenced by hypermethylation in eBL derived cell lines (4). EBV infection plays therefore a direct role in eBL pathogenesis, by altering the cellular epigenome and deregulating the expression of genes with a key role in lymphomagenesis. Indeed, i*n vitro* infection of primary B cells with EBV led to a rapid silencing of KDM2B expression. Low KDM2B mRNA level at early stages of EBV infection of primary B cells was also observed in a data set from a RNA expression profiling performed in an independent study (Manet E. personal communication). By using different means we also show that downregulation of KDM2B expression in EBV infected cells is mediated by DNMT1 recruitment onto its gene and by DNA methylation. It has already been reported that EBV can alter DNMT1 activity through its viral proteins (24). In particular Chia-Lung Tsa and colleagues showed that LMP1 activates DNMT1 activity (12) and this event requires activation of the JNK/AP1 pathway.

Here we show that cells stably expressing LMP1 display lower levels of KDM2B. The ability of LMP1 to downregulate KDM2B depends upon its NF-kB activity, as shown by using an LMP1 molecule mutated at its NF-kB activating domain, and by blocking NF-kB pathway *via* a specific chemical inhibitor. By contrast, the use of a LMP1 mutant lacking JNK pathway activation potential or a specific JNK inhibitor had no effect on KDM2B mRNA expression level. Moreover, our ChIP experiments showed that expression of LMP1 in the EBV (-) RPMI cells is able to trigger DNMT1 recruitment on the KDM2B gene. The finding that the recruitment of DNMT1 on the KDM2B gene can be hampered by treating LMP1-RPMI cells with the IκBα kinase inhibitor, further confirms its dependence on LMP1 ability to induce NF-kB pathway. Recent published data showed that HPV16 E6/E7 transforming proteins inhibit the expression of mir146-5p, known to target KDM2B transcript, which results in the increase of KDM2B expression level, in HPV16 infected cells. In contrast to E6/E7, EBV transforming protein LMP1 induces mir146-5p, *via* its ability to activate NF-kB (25). This event could contribute to the reduction of KDM2B mRNA level in cells expressing LMP1. It is therefore possible that LMP1 uses alternative mechanisms to target KDM2B: (i) increasing its DNMT1-mediated gene methylation and (ii) controlling its mRNA expression level by a specific miRNA. Further studies are needed to assess the contributions of deregulated mir146-5p levels in the events described here.

Our and other laboratories’ previous studies showed that EBV can modulate the level and the activity of different epigenetic enzymes (14, 15, 26), which in turn play a role in regulating viral genes expression. One example is provided by EZH2, whose intracellular expression level is induced in B-cells by EBV infection in a LMP1-dependent manner (27). EZH2, in turn, is recruited to the viral genome where it participates to the establishment and maintenance of EBV latency by methylating the H3 K27 in proximity to the BZLF1 and BRFL1 promoters (16). Our observation that EBV infection alters KDM2B expression prompted us to assess whether the epigenetic enzyme could regulate EBV infection and/or life cycle. Altering KDM2B expression in B-cells, prior to EBV infection, or in latently EBV-infected B cells, led to deregulated expression of all analyzed viral genes. This was consistent with the recruitment of KDM2B on their respective promoters as observed by ChIP experiments. In line with its ability to de-methylate the active chromatin mark and repress transcription, KDM2B depletion in B cells infected with EBV led to increased viral gene expression, while its forced ectopic expression had the opposite effect and caused a strong reduction of the levels of viral transcripts. Taken together, our data indicate that KDM2B plays a role in controlling EBV gene expression, for instance during the establishment of latency. Recruitment of KDM2B onto the viral episome during the first step of infection could be necessary to the repression of viral gene transcription especially at the end of the pre-latent phase when the EBV lytic genes are silenced to allow the virus to persist in resting peripheral B-cells (27). By contrast, KDM2B downregulation during the early stages of EBV infection would allow efficient viral genes expression during the pre-latent phase. Future studies will be needed to investigate the exact role of EBV-mediated KDM2B deregulation in EBV life cycle control.

It is known that in order to escape the immune-surveillance of the host and establish a chronic infection, EBV has evolved different mechanisms to maintain B-cells in a status of long -lived circulating memory B-cells and preventing them from differentiating into antibody-secreting plasma cells. A recent study has shown that EBNA3A and EBNA3C block terminal differentiation of memory B cells to plasma cells by epigenetically repressing the gene encoding for BLIMP-1, a master regulator in B cells differentiation (28). Notably a recent work has shown that KDM2B plays a key role in hematological stem cells differentiation (9). Therefore, EBV-mediated deregulation of KDM2B expression could contribute to the mechanism that prevents the cells from undergoing terminal differentiated. This hypothesis is supported by our data of a RNA profile analysis conducted on cells transfected with increasing levels of KDM2B-expressing construct. Cells harboring high KDM2B expression level showed deregulated expression of genes enriched in TNFR2 pathway, which is known to play a role in differentiation from B cells to plasma cells (19). TNFR2 also regulates the interferon pathway, an important mediator of antiviral response. Ablation of KDM2B in hematopoietic cells has been previously shown to downregulate the interferon response (9). Downregulation of KDM2B expression level upon EBV infection could therefore contribute to the virus escape from the immune-system surveillance.

Taking together our data show a novel interplay between EBV infection and the host epigenome. EBV alters KDM2B levels *via* an epigenetic mechanism involving LMP1. The histone modifier in turn, plays a role in regulating the expression of the viral and host cells genes. These data, in addition to the observed deregulated KDM2B levels in BL derived cell line, indicate that altered levels of this epigenetic enzyme could contribute to B cells transformation. Future studies aimed at investigating the functional importance of KDM2B gene methylation and downregulated expression during EBV-mediated lymphomagenic process are warranted.

## Materials and Methods

### Cell culture and treatment

Peripheral B cells were purified from blood samples using the RosetteSep human enrichment kit (Stemcell Technologies; 15064). Lymphoblastoid cell lines (LCLs) were generated in this study by infection of primary B cells from different donors, as described previously (ref). The myeloma-derived RPMI-8226 cells (http://web.expasy.org/cellosaurus/CVCL_0014) and the Burkitt lymphomas cell lines (BL), including the BL EBV(-) cell line Louckes (http://web.expasy.org/cellosaurus/CVCL_8259), were obtained from the IARC biobank. Primary and immortalized B cells were cultured in RPMI 1640 medium (GIBCO; Invitrogen life Technologies, Cergy-Pontoise, France) supplemented with 10% FBS, 100 U/ml penicillin G, 100 mg/ml streptomycin, 2 mM L-glutamine, and 1 mM sodium pyruvate (PAA, Pasching, Austria) or in advanced RPMI 1640 (LIFE TECHNOLOGIES; 12633012). EBV (Akata strain) particles produced by culturing Hone-1 EBV cells were used to infect B cells. EBV infections of B cells were performed either using recombinant WT EBV-GFP genome or using a EBV mutated strains, such as EBV_ΔLMP-1_ strain, lacking the entire LMP-1 gene. The percentage of GFP positive cells was assessed by fluorescence-activated cell sorter (FACS CANTO-Becton Dickinson). Analysis of cell cycle and apoptosis (subG1) was performed by ethanol fixing the cells and staining their DNA with propidium iodide (PI) at a final concentration of 5 µg/ml. Subsequently, cells were analyzed by FACS.

To block DNA methylation cells were treated with 5-Aza-2’-deoxycytidine ≥97%, (Sigma Aldrich; A3656) at the final concentration of 10 µM for 48 or 96h. To inhibit the different pathways, cells were treated with the IκBα kinase inhibitor Bay11-7082 (Calbiochem) to the final concentration of 10 µM, or with JNK inhibitor II SP600125 (VWR International; 420119) used at the final concentration of 10 µM. Cells were pre-incubated with the different inhibitors for 1.5h and 2h.

### Quantitative PCRs

Total RNA was extracted using the AllPrep DNA/RNA Mini Kit (Qiagen). RNA reverse transcription to cDNA was carried out by RevertAid H Minus M-MuLV Reverse Transcriptase (ThermoFischer Scientific), according to the manufacturer’s protocol. Quantitative PCR (qPCR) was performed using the MesaGreen qPCR MasterMix Plus for SYBR Assay (Eurogentec). For each primer set the qPCR was performed in duplicate and the mRNA levels obtained were normalized on the average mRNA levels of three housekeeping genes (β-globine, β-actine, GAPDH), measured in the same samples. In order to measure EBV genome copy number per cell, total DNA was extracted using the AllPrep DNA/RNA Mini Kit (Qiagen) and measured by NanoDrop. Similar amounts of DNA were used as a template for the Taqman PCR, performed according to protocol described in Accardi et al. (29). The PCR primer sequences are indicated in Table 1.

**Table 1.**
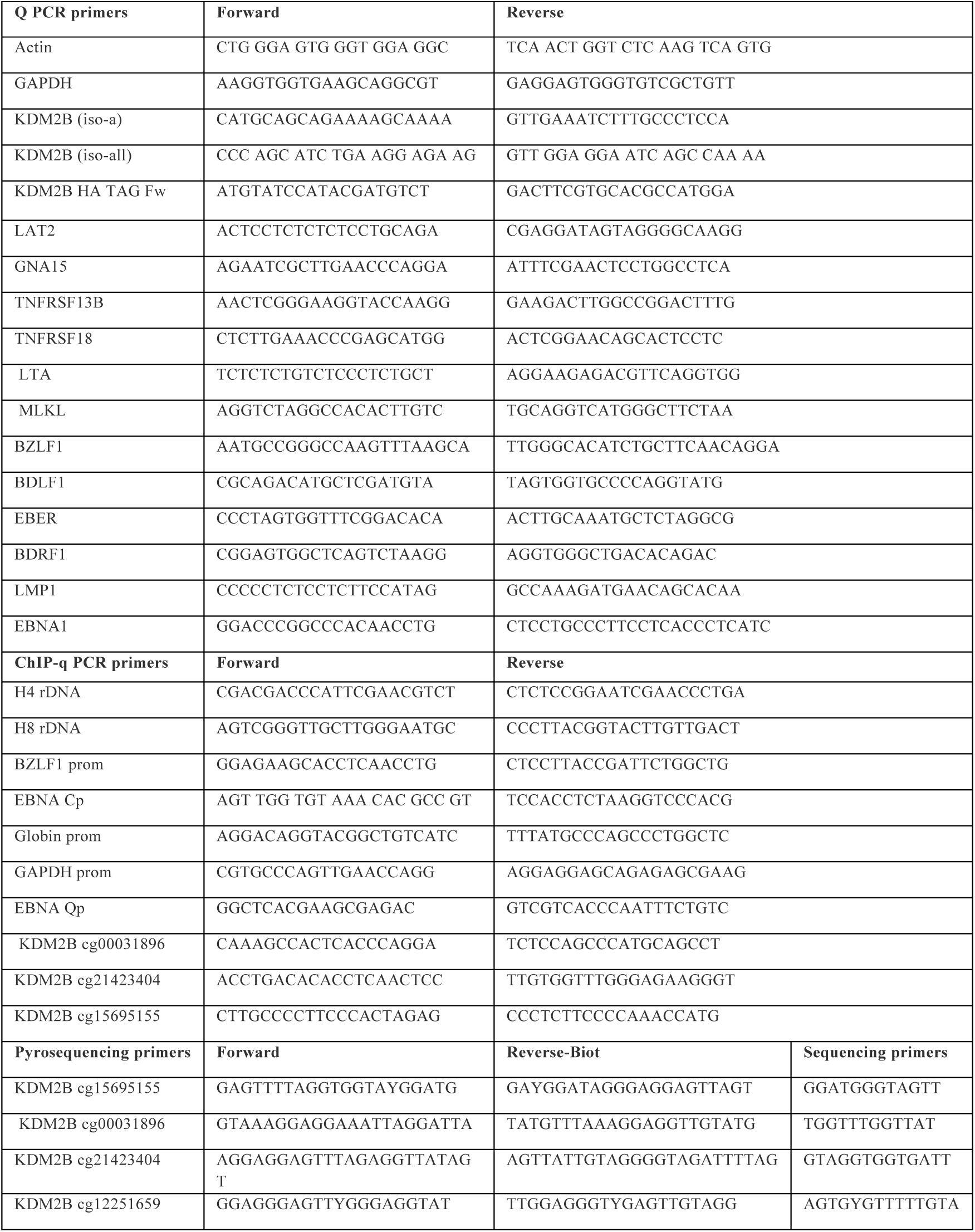
Primers used for qPCR, ChIP-PCR, and pyrosequencing are listed.

### Kdm2b overexpression and Gene-expression silencing

The KDM2B coding region was cloned into pCDNA3 vector in frame with an HA-tag at the N terminus. LCLs (1×10^7^) and Louckes (5×10^6^) were transfected with growing concentration of HA-KDM2B pCDNA3 (0,5-1,0-1,5ug) or with pCDNA3 vector as control by electroporation using the Neon Transfection System (10 µl tips) (pulse voltage, 1350V; pulse width, 30ms,pulse number 1). At 24h post-transfection, the cells were collected and processed for RNA/DNA extraction. Gene silencing of KDM2B was performed using KDM2B (human) unique 27mer siRNA duplexes (HSS150072 ThermoFischer Scientific). LCLs (1×10^7^) and Loucks (5×10^6^) were transfected with the siRNA (final concentration, 250nM) by electroporation using the Neon Transfection System (10 µl tips) (pulse voltage, 1350V; pulse width, 30ms,pulse number 1). At 48h post-transfection, the cells were collected and processed for RNA/DNA extraction. The levels of silencing were evaluated by qPCR using KDM2B specific primers indicated in Table 1.

### Immunoblotting and antibodies

Whole-cell lysate extracts were performed using lysis buffer previously described (30). Cell extracts were then fractionated by Sodium dodecyl sulfate polyacrylamide gel electrophoresis (SDS-PAGE) and processed for immunoblotting (IB) using standard techniques. The following antibodies were used for IB: KDM2B (Merck Millipore 09-864 and Abcam ab5199) and β-actin (clone C4; MP Biomedicals) and GAPDH antibody. Images were produced using the ChemiDoc XRS imaging system (Bio-Rad).

### Immuno-Fish

Fifty thousand cells were re-suspended in 5uL of PBS. The cells were gently spread on a microscope glass slide, air dried and fixed in 4% paraformaldehyde/PBS for 10 minutes at room temperature (RT). Fixes cells-containing slides were washed three times in PBS for 5 min, permeabilized with PBS-0.5% Triton X-100 (Sigma-Aldrich) for 15 min and then washed twice with PBS-Tween 0.05%. Slides were then soaked in methanol-0.3% H2O2 for 30 min, incubate 1h with antibody diluent (Dako, S3022), then 30 min with Image-iTTM FX signal enhancer. Slides were incubated overnight at 4°C with anti-KDM2B antibody (ab5199, Abcam) dilute to the concentration of 1µg/ml, followed by incubation with a secondary antibody anti-Goat (Elite kit Vector 5uL/mL). To amplify the signal the slides were incubated for 30 min at 37°C with ABC kit reagents according to the manufacture protocol. EBV DNA staining by FISH was performed as previously described (29), using a biotinylated probe to the EBV DNA genomic region BWRF1 (A300P.0100 DS-DISH-PROBES). The stained cells were visualized with a Fluorescent microscope with incubator (Nikon ECLIPSE).

### Immunohistochemistry and EBER-ISH

Immunohistochemistry analysis for KDM2B (Abcam; ab5199, dilution 1:200) was performed by an automated staining system (Ventana BenchMark ULTRA, Roche diagnostic, Monza-Italy) on FFPE 4 µm-thick sections. UltraView Universal Detection Kit (Ventana) using HRP multimer and DAB (as chromogen) was employed. ISH for EBER was carried out in each sample on 4 µm-thick section, as previously reported (31). A control slide, prepared from a paraffin-embedded tissue block containing metastatic nasopharyngeal carcinoma in a lymph node was used as positive control.

### Chromatin immunoprecipitation

Chromatin immunoprecipitation (ChIP) was performed with Diagenode Shearing ChIP and OneDay ChIP kits according to the manufacturer’s protocols. The following antibodies were used: KDM2B (Abcam ab5199), DNMT1 (Abnova MAB0079), and IgG (Diogenode). The eluted DNA was used as template for qPCR with primers designed on the promoter region of. The eluted DNA was used as template for qPCR. Primers for quantitative ChIP are listed in Table 1. The value of binding obtained for each antibody was calibrated on the input sample and normalized to the IgG values.

### Bisulfite modification and pyrosequencing

Samples for pyrosequencing were processed as previously described (Ref^3^). Primers are indicated in Table 1.

### Whole Genome Expression Analysis

Differential expression analysis was performed using Human HT-12 Expression BeadChips (Illumina) as previously described (4, 29). Probes with p value < 0.01, false discovery rate (FDR) < 0.05 and a fold-change of at least 1.5 were considered differentially expressed.

### Statistical analysis

Statistical significance was determined by Student T test. The p value of each experiment is indicated in the corresponding Figure legend. Error bars in the graphs represent the standard deviation.

## Acknowledgements

We are grateful to all members of the Epigenetics and infection and cancer biology Groups for their support. We are thankful to Elizabeth Page and Latifa Bouanzi for helping with the manuscript preparation and to Ester Sorrentino for helping with the immunohistochemistry. We finally thanks our funders: La Ligue Contre le Cancer (LNCC) Rhone (GR-IARC-2014-04-07-03 to RA), Oncostarter-(CLARA) (GR-IARC-2014-05-15-02 to RA), IARC Junior Award 2016 (AFEES-2016 to RA) and Plan cancer-INSERM (to RA).

## Additional Information

### Author Contributions Statement

RA, HG, ZH, RCVA wrote the main manuscript text and RACV, AJ, HHV, HG, AD, MPC, RCVA, GD, CS, MHM prepared figures and Tables. FLK, EM and ZH, reviewed the manuscript.

